# A structured-illumination miniscope for optically sectioned imaging and real-time neural decoding

**DOI:** 10.64898/2026.07.13.738146

**Authors:** Huixin Lin, Shu Wang, Yuhang Zhu, Zhaoyang Yin, Qingchun Guo, Jingfeng Zhou

**Affiliations:** Beijing Institute for Brain Research, Chinese Academy of Medical Sciences & Peking Union Medical College, Beijing 102206, China; Chinese Institute for Brain Research, Beijing 102206, China; Academy for Advanced Interdisciplinary Studies, Peking University, Beijing 100871, China; State Key Laboratory of Cognitive Neuroscience and Learning, Beijing Normal University & Chinese Institute for Brain Research, Beijing 100875, China; IDG/McGovern Institute for Brain Research, Beijing Normal University, Beijing 100875, China

**Author notes:** **For correspondence:** (QG); (JZ).

## Abstract

Single-photon miniscopes enable large-scale calcium imaging in freely behaving animals but are limited by out-of-focus background fluorescence that degrades image contrast and single-cell signal fidelity. Multiphoton approaches address this limitation but remain costly and complex. Here we introduce a lightweight (<3 g), low-cost structured illumination miniscope that achieves optical sectioning in freely behaving mice. Using HiLo imaging implemented with a simple Ronchi grating and time-multiplexed excitation, the system strongly suppresses background fluorescence while preserving the speed, field of view, and accessibility of widefield miniscopes, and supports optical-sectioned multi-plane imaging to increase neuronal yield. Using hippocampal recordings, we show enhanced region-of-interest (ROI)-based signal quality and spatial information readout, allowing simple ROI-averaged signals to approach the performance of offline algorithm-extracted signals. We further demonstrate a proof-of-principle closed-loop brain–machine interface enabled by rapid online signal extraction and real-time neural decoding. Together, these results establish structured illumination as a practical and accessible strategy for achieving high-contrast calcium imaging with miniaturized microscopes.

## Introduction

Miniaturized microscopes (miniscopes) are now widely used for large-scale calcium imaging in freely behaving animals, enabling population-level recordings that are difficult to obtain with conventional optical systems (***Flusberg et al., 2008; Ghosh et al., 2011; Cai et al., 2016; Barbera et al., 2016; Zong et al., 2017; Skocek et al., 2018; Zong et al., 2021, 2022; Madruga et al., 2025; Madangopal et al., 2025; Qian et al., 2026***). Single-photon widefield (WF) miniscopes are especially popular because they are lightweight, inexpensive, and easy to deploy across a range of experimental settings (***Ziv and Ghosh, 2015***). A key limitation of these systems, however, is the lack of intrinsic optical sectioning, which leads to out-of-focus background fluorescence that degrades image contrast and limits the fidelity of single-neuron signals. Computational approaches can partially mitigate these effects but operate post hoc, remain constrained by the quality of the acquired images, and can be computationally demanding, limiting their use in experiments requiring rapid online signal extraction (***Mukamel et al., 2009; Pnevmatikakis et al., 2016; Pachitariu et al., 2016; Zhou et al., 2018; Lu et al., 2018; Sità et al., 2022; Bao and Gong, 2024***). Multiphoton microscopy provides optical sectioning at the source (***Helmchen and Denk, 2005; Svoboda and Yasuda, 2006; Grienberger et al., 2022***), but its cost and technical complexity restrict broader use.

Consequently, many experiments would benefit from improved contrast and background rejection at acquisition, without sacrificing the simplicity, portability, and accessibility that have driven the widespread adoption of single-photon miniscopes. This need is especially important for real-time neural decoding and closed-loop experiments, where signals must be extracted rapidly during behavior and cannot rely on computationally intensive offline demixing. Structured illumination microscopy (SIM) offers optical sectioning by using patterned excitation to separate in-focus from out-of-focus signals (***Neil et al., 1997; Keller et al., 2010; Chen et al., 2023; Temma et al., 2024; Breuninger et al., 2007; Kalchmair et al., 2010***), without requiring multiphoton excitation or point-scanning. Among SIM approaches, HiLo microscopy (***Lim et al., 2008; Mertz and Kim, 2010; Mertz, 2011; Ford et al., 2012***) is particularly well suited for miniaturized implementations, as it requires only two images per frame—one acquired under uniform illumination and one under structured illumination—enabling efficient reconstruction with modest hardware modification and limited computational overhead.

Here we introduce a lightweight (<3 g), low-cost miniscope that implements HiLo structured illumination for calcium imaging in freely behaving mice. Using a simple Ronchi grating and dual-LED excitation, the system reduces background fluorescence at acquisition while preserving the field of view, frame rate, and usability of conventional widefield miniscopes, and supports multiplane imaging to increase neuronal yield. Using *in vivo* hippocampal recordings, we show that HiLo substantially improves ROI-based signal quality and spatial information readout, allowing simple ROI-averaged signals to approach the performance of offline algorithm-extracted signals. Finally, we demonstrate a proof-of-principle closed-loop brain–machine interface, highlighting the potential of HiLo-based miniscopes for rapid online signal extraction, real-time neural decoding, and feedback experiments.

## Results

### Implementation of a HiLo miniscope for optical sectioning

To address the challenge of background fluorescence in widefield calcium imaging, we developed a miniaturized microscope that integrated optical sectioning via structured illumination (***Figure 1***A and B). This approach used patterned illumination to selectively modulate in-focus signals while suppressing out-of-focus background fluorescence. Because out-of-focus fluorescence is only weakly modulated by the illumination pattern, HiLo reconstruction can preferentially retain the in-focus signal and reject diffuse background, thereby generating an optically sectioned image rather than simply enhancing contrast. The device was based on the principles of HiLo microscopy and employed dual illumination to acquire two complementary images using a CMOS camera: a uniformly illuminated widefield image (15 frames/sec) and a structured illumination image (15 frames/sec) generated by a Ronchi grating (***Figure 1***C and D). These paired images were combined to reconstruct a final optically sectioned image (15 frames/sec) that preserved both high- and low-spatial-frequency components of the in-focus signal, resulting in substantially enhanced contrast.

**Figure 1.**
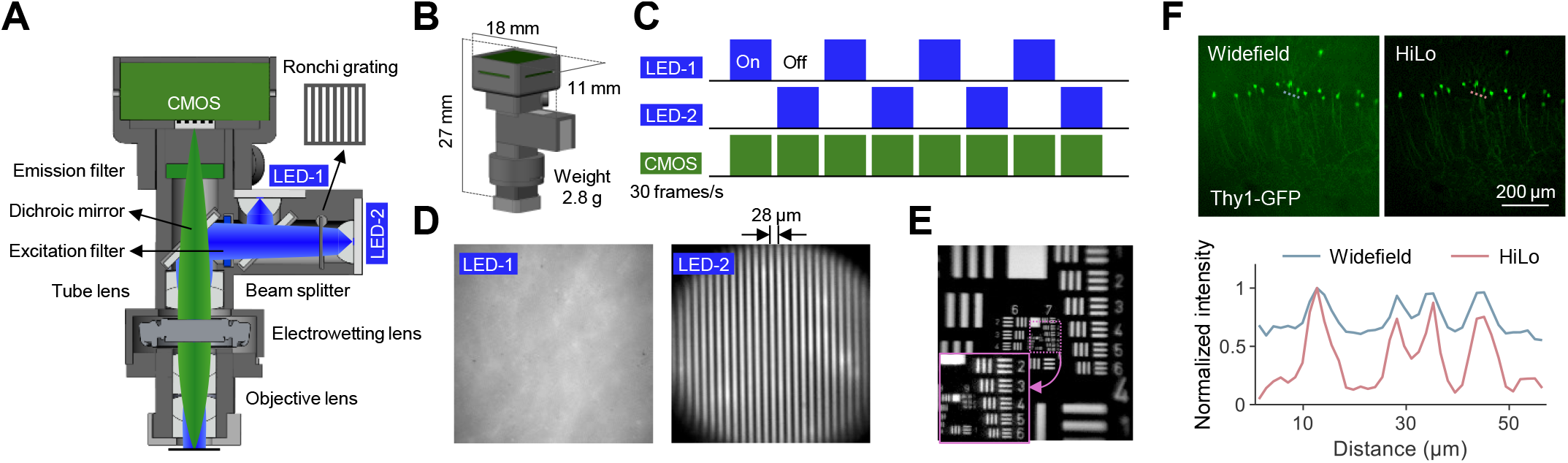
Design and performance of the HiLo miniscope. (A) Cross-sectional schematic of the HiLo miniscope. (B) Dimensions of the HiLo miniscope. (C) Time-division multiplexing scheme used to acquire uniformly illuminated and structured illumination images. The total acquisition frame rate is 30 frames per second. (D) Uniformly illuminated image and corresponding structured illumination image of a thin, uniform fluorescent plane. The Ronchi grating has a spatial period of approximately 28 *µ*m at the sample plane. (E) Uniformly illuminated image of a USAF 1951 resolution target. Inset shows a magnified view of the highlighted region; the finest resolvable feature corresponds to 256 lp/mm (Group 8, Element 1). (F) Comparison of uniformly illuminated widefield imaging and optical-sectioned HiLo miniscope imaging in a Thy1-GFP mouse brain slice. Intensity profiles along the blue and magenta line cuts illustrate the reduction of background fluorescence and the enhancement of image contrast achieved by the HiLo miniscope. **Figure 1—figure supplement 1**. Lateral and axial resolution of the HiLo miniscope.

System performance was first evaluated using a standardized optical resolution target (#55– 622, Edmund Optics), which demonstrated a lateral resolution of 256 line pairs per mm (Group 8, Element 1; ***Figure 1***E, ***Figure 1—figure Supplement 1***A and B). We further quantified the optical-sectioning capability of the system using a thin fluorescent plane and obtained an axial full width at half maximum of 23 *µ*m (***Figure 1—figure Supplement 1***C). To assess performance in biological tissue, we imaged brain slices from a Thy1-GFP mouse and observed a marked reduction in background fluorescence in HiLo-reconstructed images compared with time-multiplexed, frame-matched widefield images (***Figure 1***F). These results demonstrated that the HiLo miniscope effectively provided optical sectioning, suppressed out-of-focus background signal, and improved image contrast, supporting its utility for high-quality neural activity recordings in densely labeled preparations.

### Performance evaluation of the HiLo miniscope

We next systematically evaluated the performance of the HiLo miniscope using simulated data and *in vivo* calcium imaging in freely moving mice.

To generate simulated data, each image frame was constructed by combining three distinct components: an in-focus neural activity signal, a spatially localized out-of-focus background signal, and a global background signal (***Figure 2***A). The final widefield image was synthesized as a weighted sum of these three components. The spatial profiles of the neural activity and local background were modeled as Gaussian signals with different spatial scales to reflect their distinct contributions. The temporal dynamics of neural activity and local background across frames were derived from real hippocampal imaging data acquired with a widefield miniscope. This simulation allowed us to directly compare the recovered fluorescence signals with known ground-truth neural activity while independently modeling local and global background contamination.

**Figure 2.**
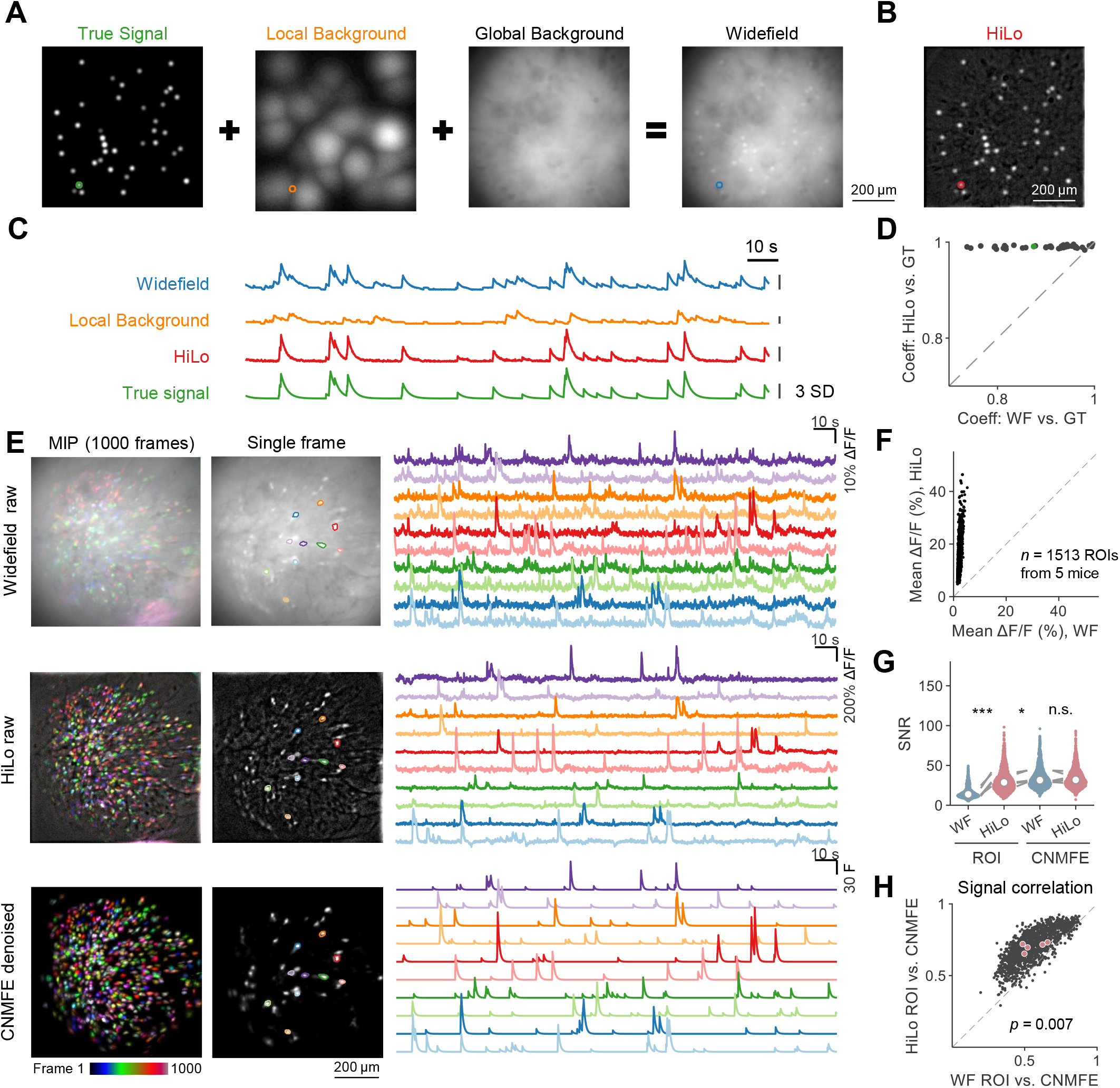
HiLo imaging improves signal fidelity and ROI-based calcium signal quality. (A) Schematic of simulated data generation. Each synthetic widefield frame was constructed as a weighted sum of three components: an in-focus neural activity signal (“true signal”), a spatially localized out-of-focus background (“local background”), and a global background signal (“global background”). (B) HiLo reconstruction of the simulated widefield image shown in (A). (C) Example fluorescence traces extracted from the simulated ROI indicated in (A) and (B). (D) Quantitative comparison of signal fidelity in simulations. Scatter plot shows the correlation coefficient between extracted signals and ground-truth activity for widefield imaging versus HiLo imaging. Each dot represents one simulated ROI; the green dot indicates the example ROI shown in (C). (E) *In vivo* calcium imaging from hippocampal CA1 in freely moving mice. Rows show raw widefield images, raw HiLo images, and CNMF-E-denoised signals. Left, maximum-intensity projections (MIPs) over 1000 frames; middle, representative single frames with example ROIs outlined; right, representative fluorescence traces from the same color-coded ROIs. (F) Comparison of mean Δ*F* ⁄*F* between widefield and HiLo ROI signals across all ROIs (*n* = 1,513 ROIs from 5 mice). Each point represents one ROI. (G) Signal-to-noise ratio comparison across extraction methods, including ROI pixel-averaged signals from widefield and HiLo images and CNMF-E–extracted signals from widefield and HiLo data. Colored dots represent individual ROIs; white dots indicate medians; lines connect session-averaged values from individual mice. (H) Similarity between ROI-based and CNMF-E–extracted signals. Scatter plot shows the correlation between ROI-derived fluorescence traces from widefield or HiLo images and denoised CNMF-E fluorescence traces extracted from the corresponding widefield dataset. Black dots represent individual ROIs, and colored dots indicate session means. Statistical significance in (G and H) was assessed using paired *t*-tests on mouse-level mean values (*n* = 5 mice; \**p* < 0.05, \*\*\**p* < 0.001; n.s., not significant). **Figure 2—video 1**. Representative HiLo calcium imaging movie.

We then processed the simulated data using the same reconstruction algorithm employed by the HiLo miniscope to generate the final HiLo images (***Figure 2***B). Comparison of the simulated widefield and HiLo images revealed a substantial reduction in both local and global background fluorescence in the HiLo reconstructions, while the true neural activity signals were largely preserved (***Figure 2***C). Direct comparison of correlations between the widefield and HiLo signals relative to the ground-truth activity showed that the HiLo method recovered neural signals with markedly higher fidelity than conventional widefield imaging (***Figure 2***D). These simulations therefore confirmed that HiLo reconstruction can suppress background contamination while preserving in-focus neural activity.

We then evaluated HiLo performance using *in vivo* calcium imaging in freely moving mice. The HiLo miniscope was mounted on the animals’ heads to record activity from the hippocampal CA1 region through a gradient-index (GRIN) lens. Uniformly illuminated widefield images and structured-illumination images were acquired in an alternating, time-multiplexed manner to enable direct comparison. Representative ROI pixel-averaged signals showed a pronounced reduction in back-ground fluorescence and a corresponding improvement in signal-to-noise ratio (SNR) in HiLo images relative to widefield images (***Figure 2***E, ***Figure 2—video 1***). Across multiple imaging sessions and animals, HiLo imaging consistently outperformed widefield imaging in terms of background suppression and SNR for ROI-based signals (WF, ROI vs. HiLo, ROI: *p* = 0.0006; paired *t*-test; ***Figure 2***F and G).

In contrast, after offline CNMF-E extraction, SNRs were not significantly different between wide-field and HiLo data (WF, CNMF-E vs. HiLo, CNMF-E: *p* = 0.087; paired *t*-test; ***Figure 2***G), indicating that offline source extraction reduced the modality-dependent difference. Importantly, simple pixel-averaged HiLo ROI signals approached the SNRs obtained with offline CNMF-E extraction, although they remained modestly lower than WF CNMF-E signals (HiLo, ROI vs. WF, CNMF-E: *p* = 0.043; paired *t*-test; ***Figure 2***G). Consistent with this observation, HiLo ROI signals showed significantly higher correlations with WF CNMF-E–denoised signals than did WF ROI signals (*p* = 0.007; paired *t*-test; ***Figure 2***H), indicating that HiLo ROI measurements more closely resembled the denoised neural activity patterns recovered by CNMF-E extraction. Thus, the principal advantage of HiLo imaging was most evident before offline demixing—acquisition-level background suppression substantially improved simple ROI-based signals, bringing them close to the quality of algorithm-extracted signals. This distinction is important because CNMF-E can selectively recover activity components from background-contaminated images, whereas HiLo improves the optical image itself by suppressing out-of-focus fluorescence at acquisition.

### Optical-sectioned multi-plane imaging increases neuronal yield

To test whether the HiLo miniscope could support multi-plane or volumetric imaging, we combined acquisition-level optical sectioning by HiLo background suppression with voltage-controlled focal-depth modulation through the electrowetting liquid lens already integrated into the miniscope design (***Figure 1***A). We first calibrated the relationship between applied voltage and axial focal position, confirming focal-depth modulation across an axial range of up to 450 *µ*m (***Figure 3—figure Supplement 1***A). We then validated this capability in Thy1-GFP mouse brain slices, where sequential imaging across focal depths produced high-contrast optical-sectioned image stacks spanning up to 132 *µ*m (***Figure 3***A, ***Figure 3—figure Supplement 1***B). These results showed that the liquid-lens-based HiLo miniscope could perform optical-sectioned multi-depth imaging, thereby establishing the technical basis for multi-plane recordings *in vivo*.

**Figure 3.**
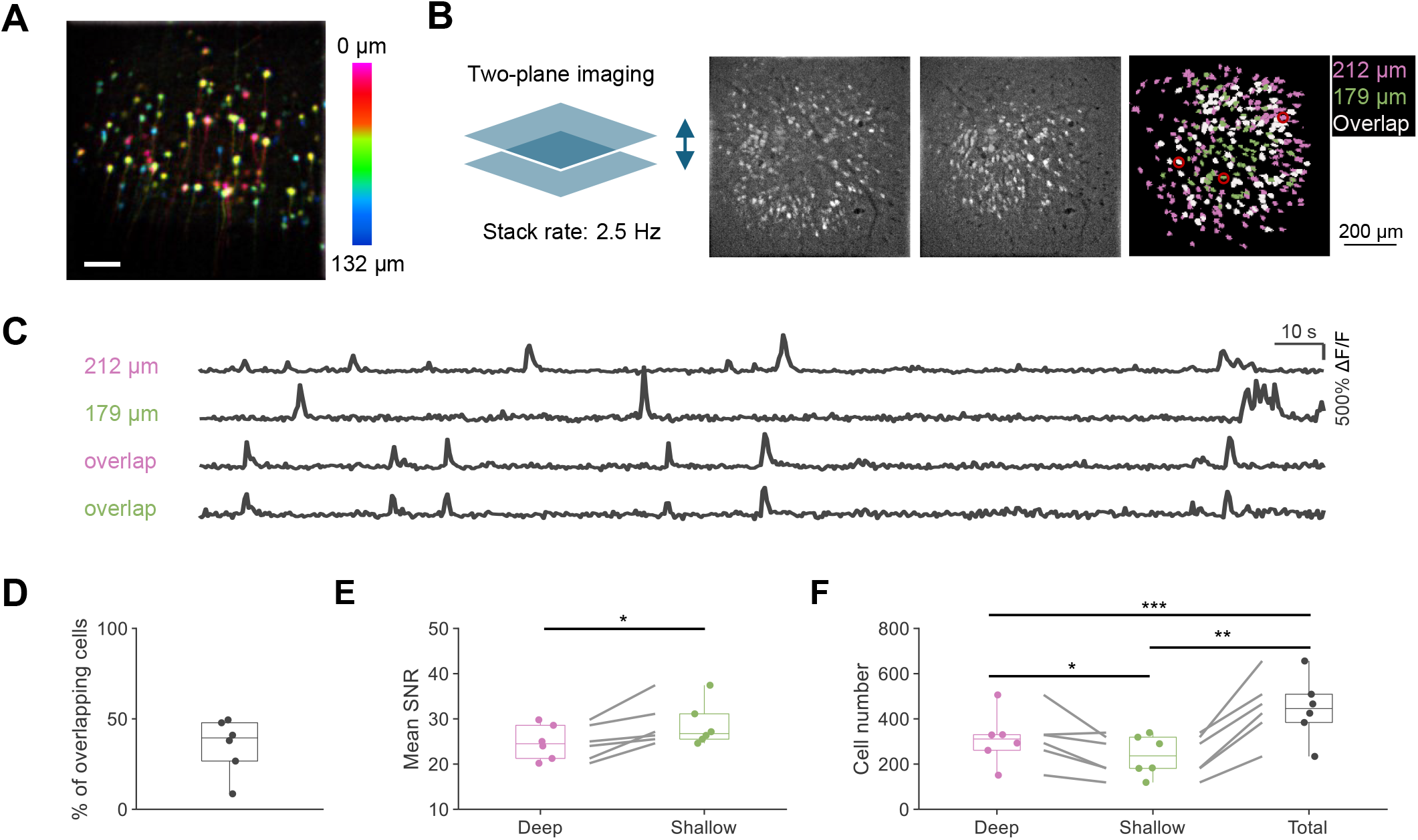
Optical-sectioned multi-plane imaging increases the yield of imaged neurons. (A) Color-coded composite image sampled from a Thy1-GFP mouse brain slice across multiple focal depths, illustrating volumetric coverage enabled by optical-sectioned HiLo imaging. Color indicates relative imaging depth. Scale bar, 100 *µ*m. (B) Schematic and representative maximum-intensity projection (MIP) images illustrating *in vivo* two-plane imaging. Example fluorescence images acquired at depths of 179 *µ*m and 212 *µ*m are shown, together with detected neurons color-coded by imaging plane and overlap (magenta, 212 *µ*m; green, 179 *µ*m; white, neurons detected in both planes). (C) Representative calcium activity traces (Δ*F* ⁄*F*) from neurons detected exclusively in the deeper plane (212 *µ*m), exclusively in the shallower plane (179 *µ*m), and in both planes (overlap), demonstrating reliable signal extraction across imaging depths. (D) Percentage of neurons detected in both imaging planes relative to the total number of neurons identified across planes. (E) Comparison of mean signal-to-noise ratio (SNR) between neurons detected in the deeper and shallower imaging planes. (F) Number of neurons detected in the deeper plane, the shallower plane, and after combining neurons from both planes. In (D)–(F), each dot represents one imaging session. In (E) and (F), paired lines indicate within-session comparisons. Boxes represent the median and interquartile range, and whiskers indicate the minimum and maximum values. Statistical significance was assessed using paired *t*-tests (*n* = 6 mice; \**p* < 0.05, \*\**p* < 0.01, \*\*\**p* < 0.001). **Figure 3—figure supplement 1**. Multi-plane imaging with a liquid-lens–based HiLo miniscope.

For *in vivo* hippocampal recordings, we employed a two-plane configuration with the focal planes separated by 33 *µ*m and alternated between planes at an imaging rate of 2.5 Hz (***Figure 3***B). Representative calcium traces extracted from neurons detected exclusively in each plane, as well as from neurons detected in both planes, exhibited clear and well-resolved activity patterns (***Figure 3***C), demonstrating reliable signal extraction across imaging depths under optical-sectioned multi-plane imaging conditions.

Given the high density of pyramidal neurons in dorsal CA1, a subset of neurons was detected in both imaging planes. Quantification of this overlap showed that overlapping neurons constituted a minority of the total population imaged across planes (***Figure 3***D), indicating that each imaging plane contributed a largely distinct set of neurons. Signal quality differed modestly across depth. Neurons detected in the shallower plane exhibited higher mean SNR than those detected in the deeper plane, although robust calcium signals were obtained from both planes (deep vs. shallow: *p* = 0.013; paired *t*-test; ***Figure 3***E). Combining neurons from both planes substantially increased the total number of neurons recorded relative to either plane alone, yielding a larger overall population for downstream analysis (deep vs. shallow: *p* = 0.049; shallow vs. total: *p* = 0.001; deep vs. total: *p* = 0.0004; paired *t*-tests; ***Figure 3***F).

Together, these results demonstrate that the HiLo miniscope not only improves image contrast and ROI signal quality through optical background suppression, but also leverages its integrated electrowetting liquid lens to perform optical-sectioned multi-plane imaging. This combination increases neuronal yield while maintaining reliable signal quality across imaging depths.

### HiLo miniscope enhances ROI-based spatial readout from the hippocampus

Given the improved performance of HiLo imaging in suppressing background fluorescence for ROI-based signals, we next examined whether this acquisition-level improvement enhanced the read-out of behaviorally relevant neural activity. We recorded calcium activity from dorsal hippocampal CA1 while mice freely explored an open-field arena (***Figure 4—figure Supplement 1***) and compared the spatial information content extracted from widefield and HiLo images.

**Figure 4.**
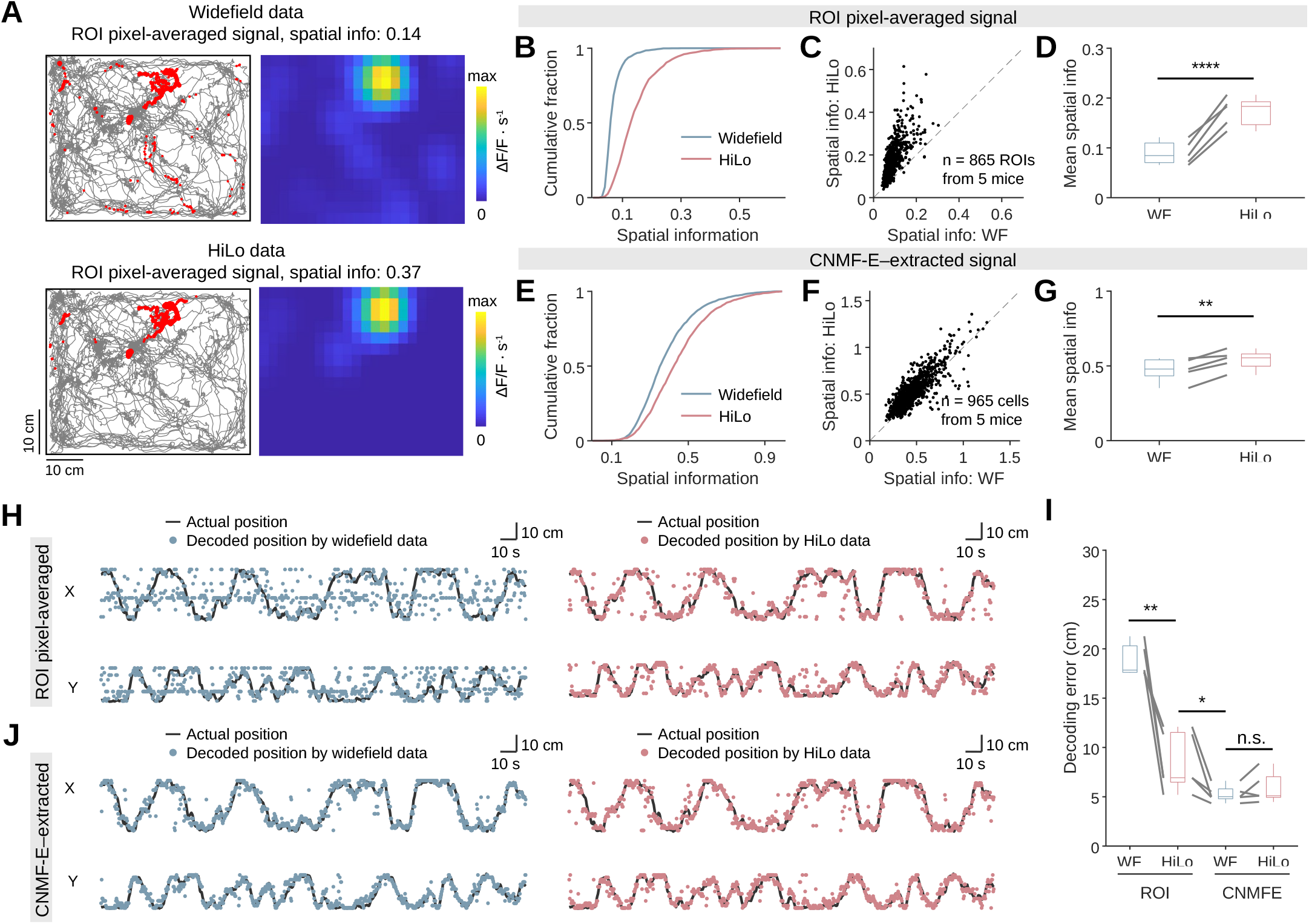
HiLo imaging enhances ROI-based spatial readout from hippocampal CA1. (A) Representative matched ROI from dorsal hippocampal CA1. Left: animal trajectory (gray) overlaid with calcium event locations (red); only events exceeding 3 standard deviations of Δ*F* ⁄*F* are shown. Right: corresponding spatial tuning map, computed with 2.5 × 2.5 cm spatial bins and smoothed with a Gaussian kernel (σ = 3.5 cm). (B) Cumulative distributions of spatial information for individually extracted ROI pixel-averaged signals from widefield and HiLo images. (C) Comparison of spatial information for matched ROIs identified in both widefield and HiLo images; each dot represents one ROI. (D) Summary of mean spatial information for matched ROIs across imaging modalities. (E)–(G) Same analyses as in (B)–(D), performed on signals extracted using the CNMF-E algorithm. (H) Bayesian decoding of animal position using ROI pixel-averaged signals from widefield and HiLo images. Actual trajectories are shown in black, and decoded positions are overlaid as colored dots for each imaging modality. (I) Decoding performance quantified as median decoding error. (J) Same decoding analyses as in (H) using CNMF-E–extracted signals. In (D), (G), and (I), paired lines indicate within-session comparisons; boxes show median and interquartile range; whiskers indicate extreme values; statistical significance was assessed using paired *t*-tests (*n* = 5 mice; \**p* < 0.05, \*\**p* < 0.01, \*\*\*\**p* < 0.0001; n.s., not significant). **Figure 4—figure supplement 1**. HiLo calcium imaging in an open-field arena. **Figure 4—figure supplement 2**. Spatial information readout from hippocampal CA1 during free exploration.

For direct comparison between imaging modalities, we applied the spatial footprints of putative neurons identified by CNMF-E to the raw images and computed ROI pixel-averaged signals for spatial information analysis. This analysis was designed to isolate the effect of acquisition-level background suppression on simple ROI-based signals, without requiring offline demixing of the fluorescence traces. Representative spatial tuning maps from a matched ROI showed markedly reduced activity outside the place field in HiLo images compared to widefield images (***Figure 4***A, ***Figure 4—figure Supplement 2***A). Consistently, ROI pixel-averaged signals derived from HiLo images conveyed significantly higher spatial information than those from widefield images, both for individually extracted ROIs (***Figure 4***B) and for matched ROIs identified across the two modalities (WF vs. HiLo: *p* < 0.0001; paired *t*-test; ***Figure 4***C and D). These results indicate that optical background suppression improved the behavioral specificity of minimally processed ROI signals.

For CNMF-E–extracted fluorescence signals, spatial information was also higher in HiLo data than in widefield data, but the magnitude of this improvement was smaller (WF vs. HiLo: *p* = 0.008; paired *t*-test; ***Figure 4***E–G). Similar results were obtained using deconvolved CNMF-E signals, for which spatial information was also higher in the HiLo dataset (*p* = 0.009; paired *t*-test;***Figure 4— figure Supplement 2***C). Because spatial information measurements are sensitive to baseline activity levels, with sparser activity often yielding higher spatial information values, we further examined whether differences in activity rate contributed to the observed effect. Consistent with the enhanced background suppression provided by HiLo imaging, deconvolved HiLo signals exhibited lower event rates than widefield signals (*p* < 0.0001; paired *t*-test; ***Figure 4—figure Supplement 2***B). To control for this difference, we randomly removed spike events from the widefield dataset to match the activity level observed in the HiLo data and repeated the spatial information analysis. After activity matching, spatial information no longer differed between the two datasets (*p* = 0.083; paired *t*-test; ***Figure 4—figure Supplement 2***D–E). Thus, HiLo improved spatial information read-out most strongly when neural activity was extracted using simple ROI averaging, whereas offline source extraction reduced the performance gap between widefield and HiLo imaging.

To assess spatial information at the population level, we trained Bayesian decoders (***Zhang et al., 1998; Davidson et al., 2009; Lai et al., 2023; Ziv et al., 2013***) to predict the animals’ positions using ROI pixel-averaged signals from widefield and HiLo images (***Figure 4***H). Decoding error was significantly lower when using HiLo-derived signals (WF vs. HiLo: *p* = 0.005; paired *t*-test; ***Figure 4***I). In contrast, decoders trained on CNMF-E–extracted signals showed comparable performance between widefield and HiLo data (WF vs. HiLo: *p* = 0.25; paired *t*-test; ***Figure 4***I–J, ***Figure 4—figure Supplement 2***F and G), indicating that HiLo background suppression preserves spatial information while enhancing its accessibility in raw ROI signals. Notably, decoder performance using HiLo ROI signals approached that obtained with CNMF-E–extracted signals (*p* = 0.046; paired *t*-test; ***Figure 4***I), suggesting that acquisition-level background suppression enables simple ROI measurements to capture much of the behaviorally relevant information typically recovered through computationally intensive demixing. This comparison further supports the conclusion that HiLo is especially advantageous for rapid ROI-based readout.

Together, these results demonstrate that HiLo imaging enhances spatial information readout in hippocampal recordings, particularly when neural activity is accessed through simple ROI-based measurements, highlighting its potential for rapid online signal extraction, real-time neural decoding, and behavioral analysis.

### Closed-loop brain–machine interface using rapid online ROI signal extraction

Real-time decoding is particularly valuable in closed-loop control systems, especially within BMI applications. Based on the improved quality of ROI-based HiLo signals, we extended its use to a closed-loop BMI paradigm. Our goal was to test whether HiLo-based imaging, when integrated with a real-time decoding algorithm, could support closed-loop reward control based on hippocampal population activity (***Figure 5***A, ***Figure 5—figure Supplement 1***A).

**Figure 5.**
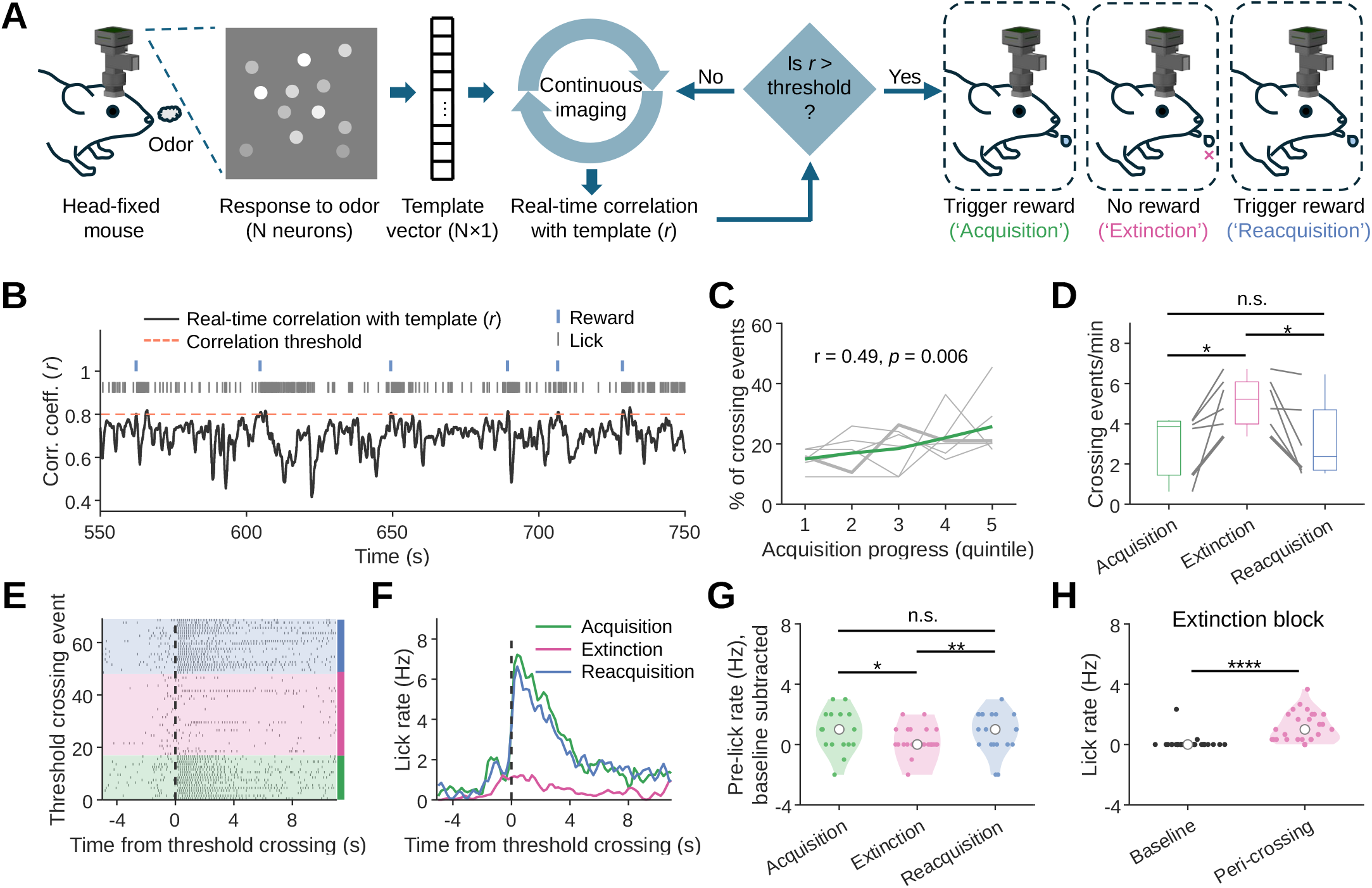
Closed-loop hippocampal BMI training. (A) Experimental setup for a closed-loop BMI paradigm. The real-time decoding algorithm compares ongoing population vector activity to a pre-established odor-evoked template. If the correlation exceeds a threshold, a reward is triggered. Training consists of acquisition, extinction, and reacquisition phases, with rewards delivered during acquisition and reacquisition but not extinction. (B) Example session of real-time correlation of population vector activity with the template during training. Red dashed line represents correlation threshold. Vertical blue lines are reward delivery and gray bars are licks. (C) Threshold-crossing events (% of total) across the acquisition process. Gray lines indicate individual animals, with the bolded gray line showing the example mouse in (B), (E)–(H); green line shows the population average. Correlation between acquisition progress and threshold-crossing events is tested using Spearman correlation. (D) Frequency of crossing events per minute during the acquisition, extinction, and reacquisition phases. Bolded gray line corresponds to the example mouse. Paired lines indicate within-session comparisons; boxes show median and interquartile range; whiskers indicate extreme values. (E) Example behavioral session showing lick rasters aligned to threshold-crossing events (dashed line); shaded regions indicate training blocks. (F) Lick rate as a function of time relative to threshold-crossing events for the example session. (G) Lick rate preceding threshold-crossing events (−2 to −1 s), corrected by subtracting baseline lick rate (−4 to −3 s), across acquisition, extinction, and reacquisition blocks for the same session. Statistical significance was assessed using rank sum test (\**p* < 0.05, \*\**p* < 0.01; n.s., not significant). (H) Baseline (−6 to −3 s) and peri-crossing (−1 to 2 s) lick rates during extinction. Statistical significance was assessed using a paired *t*-test (*n* = 32 crossing events; \*\*\*\**p* < 0.0001). In (G) and (H), colored dots represent individual crossing events; white dots indicate medians. **Figure 5—figure supplement 1**. Closed-loop calcium imaging and lick behavior during BMI training.

To this end, mice were head-fixed and presented with repeated odor stimuli while CA1 activity was recorded using the HiLo miniscope. Spatial footprints of putative neurons were rapidly identified using Suite2p and used to extract pixel-averaged ROI signals from raw HiLo images. Odor-evoked population activity was represented as a population vector (PV), which served as a template for detecting time points at which ongoing population activity resembled the odor-evoked pattern. During training, moment-to-moment PV activity was continuously compared to the template, and a sucrose reward was delivered whenever the PV correlation exceeded a predefined threshold. Each session consisted of acquisition, extinction, and reacquisition blocks, during which threshold-crossing events triggered reward delivery in the acquisition and reacquisition blocks but not during extinction (***Figure 5***A).

This closed-loop training system was successfully implemented (***Figure 5***B). During the acquisition phase, the frequency of successful threshold-crossing events increased over time (*p* = 0.006, Spearman correlation; ***Figure 5***C), consistent with adaptive modulation of hippocampal population activity to obtain reward under the closed-loop contingency. When reward delivery was omitted during extinction, the rate of threshold-crossing events increased, resembling an extinction burst (***Shahan and Avellaneda, 2025***), and subsequently decreased when reward delivery was reinstated during reacquisition (acquisition vs. extinction: *p* = 0.029; extinction vs. reacquisition: *p* = 0.033; acquisition vs. reacquisition: *p* = 0.80; paired *t*-tests; ***Figure 5***D), suggesting task-dependent modulation of neural activity.

Although the closed-loop training paradigm did not require an overt conditioned response, one mouse exhibited contingency-dependent licking behavior reminiscent of anticipatory licking in classical conditioning tasks (***Figure 5***E–H, ***Figure 5—figure Supplement 1***B). Specifically, lick rates preceding threshold-crossing events declined during extinction and recovered during reacquisition (acquisition vs. extinction: *p* = 0.017; extinction vs. reacquisition: *p* = 0.006; acquisition vs. reacquisition: *p* = 0.97; rank sum test; ***Figure 5***G), providing an additional behavioral readout of the closed-loop contingency in this animal. Moreover, peri-crossing lick rates during the extinction block were significantly elevated relative to baseline (*p* < 0.0001, paired *t*-test; ***Figure 5***H), further suggesting that threshold-crossing events were behaviorally meaningful in this example session.

These results demonstrate that HiLo microscopy enables rapid online ROI signal extraction and real-time neural decoding with sufficient signal quality for closed-loop BMI control. This capability highlights the utility of acquisition-level optical background suppression for experiments that require fast access to population activity during behavior.

## Discussion

The miniature structured illumination microscope described here addresses a central limitation of widely used single-photon miniscopes: the absence of optical sectioning at acquisition. By integrating HiLo structured illumination into a lightweight, head-mounted design, the system improves image contrast and single-cell signal fidelity while preserving the simplicity, affordability, and ease of use that have driven the broad adoption of widefield miniscope imaging. Rather than replacing existing approaches, it provides a complementary solution that bridges conventional widefield miniscopes and more complex multiphoton systems, enabling higher-quality population recordings in freely behaving animals.

A key advantage of this design is that background fluorescence is suppressed during image acquisition rather than corrected post hoc. While computational approaches can partially mitigate background contamination after acquisition (***Mukamel et al., 2009; Pnevmatikakis et al., 2016; Pachitariu et al., 2016; Zhou et al., 2018; Lu et al., 2018; Bao and Gong, 2024***), HiLo provides optical sectioning directly at the source, resulting in cleaner images before downstream processing. Our results show that this acquisition-level improvement is especially important for simple ROI-based signal extraction. HiLo substantially improved ROI-averaged calcium signals and allowed them to approach the quality of signals obtained with offline source-extraction algorithms. These findings suggest that optical sectioning can recover a substantial fraction of the information typically obtained through computational processing while preserving a simple and efficient signal-extraction workflow. This acquisition-level improvement may be especially valuable in settings where rapid access to neural activity is desirable, including real-time neural decoding and closed-loop experimental paradigms.

These gains are achieved with minimal added hardware complexity. Using a simple Ronchi grating and time-multiplexed illumination, HiLo reconstruction requires only two images per frame and modest computational processing, facilitating miniaturization and compatibility with existing open-source miniscope ecosystems. In addition, the incorporation of an electrowetting liquid lens enables optical-sectioned multi-plane imaging, increasing neuronal sampling without substantially increasing device weight or experimental overhead. Although demonstrated here at moderate frame rates, this capability supports studies focused on distributed population codes and neural representations that benefit from increased sampling across depth.

We evaluated the system using hippocampal calcium imaging as a demanding test case (***Flusberg et al., 2008; Cai et al., 2016; Skocek et al., 2018; Zong et al., 2022; Madruga et al., 2025; Ziv et al., 2013; Lin and Zhou, 2024***), given the high neuronal density and substantial background fluorescence characteristic of this region. In this context, improved contrast enabled more reliable extraction of neuronal signals and enhanced decoding of spatial information. Importantly, the improvement was most pronounced for ROI-based signals, whereas offline source extraction reduced the performance gap between widefield and HiLo imaging. This pattern indicates that the major practical benefit of HiLo is not simply higher final performance after offline processing, but improved access to useful neural signals directly from minimally processed images. The strong performance of HiLo ROI signals indicates that acquisition-level optical sectioning effectively suppresses diffuse background contamination, a major source of signal degradation in single-photon imaging. Nevertheless, HiLo does not explicitly separate overlapping neuronal sources. Consistent with this distinction, decoding performance obtained from HiLo ROI signals remained slightly below that achieved using CNMF-E-processed widefield data. Together, these findings suggest that optical sectioning can recover much of the signal quality typically achieved through computational demixing, while additional gains may still arise from algorithmic source separation. Therefore, HiLo should be viewed as complementary to, rather than a replacement for, advanced computational methods.

Because these improvements arise from acquisition-level signal quality rather than task-specific optimization, they are expected to generalize to other brain regions and experimental paradigms in which background contamination limits widefield imaging performance. As a proof of principle, we also implemented a closed-loop brain–machine interface paradigm, demonstrating that the system supports rapid online ROI signal extraction, real-time neural decoding, and feedback. In this implementation, spatial footprints were identified before closed-loop training and then used for fast ROI-based readout, avoiding the need for real-time source demixing during the experiment. These demonstrations are not intended to define the primary application domain of the device, but rather to illustrate its compatibility with experiments that place stringent demands on signal fidelity and timing.

Compared with multiphoton microscopy, including recent miniaturized two-photon systems (***Zong et al., 2017, 2022; Madruga et al., 2025; Qian et al., 2026; Ozbay et al., 2018; Klioutchnikov et al., 2020, 2023; Zhao et al., 2023b,a***), the HiLo miniscope represents a balance between optical performance and system complexity. Multiphoton approaches offer superior optical sectioning, deeper tissue penetration, and reduced sensitivity to scattering, making them well suited for imaging in dense or deep tissue (***Svoboda and Yasuda, 2006; Grienberger et al., 2022***). These advantages, however, require more complex optical architectures, higher power consumption, and tighter constraints on alignment and system stability. By contrast, the HiLo approach improves contrast and background suppression at acquisition while preserving the low power requirements and simple optical design of single-photon miniscopes. Although imaging depth remains more limited than in multiphoton systems, this design provides an effective solution for population-level calcium imaging and applications requiring rapid online signal extraction in freely behaving animals. Future refinements in illumination patterns, reconstruction efficiency, and optical components may further extend performance across experimental preparations.

The capabilities of the HiLo miniscope should be interpreted in light of several optical and computational trade-offs. First, although HiLo suppresses out-of-focus background fluorescence, it does not provide the scattering rejection or penetration depth of multiphoton microscopy. Second, HiLo improves the optical image at acquisition but does not by itself demix overlapping neuronal sources; for densely packed populations, computational source extraction may still provide additional benefits. Third, the multi-plane implementation trades imaging speed for axial sampling, and future optimization of sensor frame rate, illumination timing, and liquid-lens response time could further improve volumetric imaging performance.

In summary, this work establishes structured illumination as a practical and accessible strategy for improving image quality in miniaturized calcium imaging. By enhancing contrast at acquisition while retaining the core advantages of single-photon miniscopes and requiring only minimal hardware modification, this approach provides a broadly useful resource for systems neuroscience studies that rely on stable, high-quality population recordings during natural behavior and for closed-loop experiments that require rapid neural signal extraction and real-time feedback.

## Methods and Materials

### Animal subjects

One 12-week-old male Thy1-GFP mouse (Tg(Thy1-EGFP)MJrs/J; The Jackson Laboratory, strain #007788) was used for the brain-slice demonstration. Seven Thy1-GCaMP6f mice (five males and two females; C57BL/6J-Tg(Thy1-GCaMP6f)GP5.17Dkim/J; The Jackson Laboratory, strain #025393), aged 3–16 months at the start of experiments, were used for *in vivo* calcium imaging. Mice were group-housed (up to five animals per cage) under a 12 h light/12 h dark cycle. All experiments were conducted during the dark phase. Ambient temperature was maintained at 23–25 ^°^C with 40%– 50% relative humidity. Animals had ad libitum access to food and water until 2–3 days before closed-loop experiments, after which water restriction was implemented to maintain body weight at ≥80% of free-drinking levels. All behavioral experiments were conducted at the Chinese Institute for Brain Research (CIBR), Beijing. All animal care and experimental procedures were approved by the Institutional Animal Care and Use Committee of CIBR (AP# CIBRIACUC-037).

### Surgical procedures

Mice were anesthetized with isoflurane (5% for induction and 1–1.5% for maintenance), and ophthalmic ointment was applied to prevent eye dryness. After disinfection with 75% (v/v) alcohol, the scalp was incised to expose the skull, which was leveled by adjusting the nose bar and two ear bars. For CA1 imaging, a craniotomy was performed above the right dorsal CA1, with a 1.2 mm diameter and centered at AP −2.3 mm, ML 2.0 mm. Overlying cortical tissue was aspirated using a blunted 28-gauge needle connected to a vacuum under irrigation with sterile, cold saline until the corpus callosum became visible. A blunted 30-gauge needle was then used to remove large superficial fibers, leaving a thin fiber layer intact. Sterile Gelfoam was applied to stop bleeding. A GRIN lens (1 mm diameter, 4 mm length; GoFoton) was lowered to the tissue surface and then advanced 50 *µ*m deeper to accommodate brain swelling. Dental cement (C&B Metabond) was applied to fix the lens to the skull and attach a titanium headplate. Three weeks after lens implantation, animals were head-restrained and examined for fluorescence signals using the miniscope described in this study mounted on a baseplate. The field of view and the distance between the objective and the GRIN lens were adjusted until blood vessels and putative neuronal somas were visible near the center of the adjustable focus range. The baseplate was then cemented to the skull, and the miniscope was replaced with a 3D-printed protective cover. Animals were allowed to recover for at least three days after baseplate attachment before daily handling and subsequent behavioral procedures.

### Histology

At the end of behavioral and imaging experiments, mice were deeply anesthetized and transcardially perfused with saline, followed by 4% paraformaldehyde (PFA) in phosphate-buffered saline (PBS; pH ~7.4). Brains were carefully extracted and post-fixed in 4% PFA for at least 6 h at room temperature (25 ^°^C), then cryoprotected in 30% (w/v) sucrose in PBS until the tissue sank, typically ~12 h at room temperature. After embedding and freezing, brains were sectioned coronally at 50 or 200 *µ*m using a cryostat microtome (CM3050 S, Leica). Sections were mounted on glass slides and counterstained with DAPI. Fluorescence signals, including GFP, GCaMP6f, and DAPI, were imaged using a slide scanner (VS120, Olympus) to verify lens placement.

### Basic principle of HiLo microscopy

HiLo microscopy (***Lim et al., 2008; Mertz and Kim, 2010; Mertz, 2011; Ford et al., 2012***) is one of the implementations of optical sectioning structured illumination microscopy (OS-SIM) (***Neil et al., 1997; Keller et al., 2010; Chen et al., 2023; Breuninger et al., 2007; Kalchmair et al., 2010***), which uses patterned illumination to modulate in-focus signals and produce an optically sectioned image with out-of-focus background suppressed. In HiLo microscopy, a uniformly illuminated image 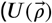, where 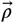 represents the 2D spatial coordinate) and a structured-illumination image 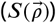 are required to recover the high-frequency and low-frequency components of the in-focus signal. They can be written respectively as (***Mertz and Kim, 2010***):

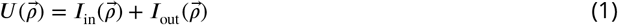

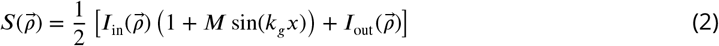

where 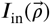 and 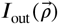 are in-focus and out-of-focus signals, respectively. In 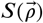, the in-focus illumination is modulated in a sinusoid pattern in the *x*-direction with frequency *k*_*g*_ and contrast *M*.

Because the high-frequency components attenuate quickly during defocusing, 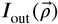 contains mainly low-frequency information, which means 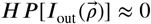 Thus, the high-frequency components of 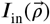 can be directly recovered by using a high-pass filter (the cutoff frequency was much smaller than *k*_*g*_) to 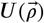, obtaining

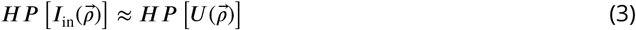

To recover the lost low-frequency components of the in-focus signal, a demodulated image is obtained firstly:

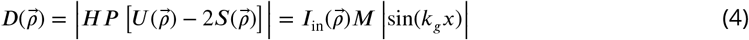

By applying a low-pass filter directly to this demodulated image, we then get

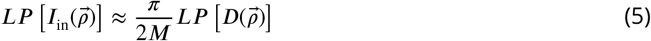

The final in-focus image 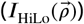 is recovered by combining Eqs. (3) and (5):

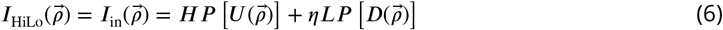

where η is a scaling factor to make the transition at full frequency bandwidth seamless.

### Implementation of the HiLo miniscope

The HiLo miniscope is designed based on the open-source solutions provided by UCLA Miniscope (www.miniscope.org) and Ninscope (***de Groot et al., 2020***). As illustrated in Figure 1A, two blue LED light sources (470 nm, LXML-PB02, Lumileds) were employed as the excitation light sources. These were collimated using half-ball lenses (3 mm diameter, #47–269, Edmund Optics) prior to their combination into a single excitation light path via a customized beam splitter (3.5 × 5 × 0.5 mm, 50:50). The excitation light passing through the excitation filter (3.5 × 3.5 × 0.5 mm, ET470/40, Chroma) was subsequently reflected by the dichroic mirror (3.5 × 5 × 0.5 mm, T495lpxr, Chroma) and focused onto the sample through the tube lens (4 mm diameter, #63-692, Edmund Optics) and the objective lens module (3 mm diameter, #45-089 × 2, Edmund Optics). The fluorescent light emitted from the sample was filtered by the emission filter (4 × 4 × 1 mm, ET525/50 m, Chroma) and relayed to generate the final image on the CMOS camera (PYTHON480, ON Semiconductor) utilizing the same lens combination. A transmissive Ronchi grating was positioned between the LED2 half-ball lens and the beam splitter to produce structured illumination. The positions of the Ronchi grating and CMOS camera were adjusted so that they were conjugate to the object plane. An electrowetting lens (2.5 mm clear aperture, A-25H0, Corning) was employed to modify the focus plane on the sample. All these optical components were ultimately assembled into a housing fabricated from black resin (FLGPBK04, Formlabs) using a 3D printer (Form 3, Formlabs). The base plate and DAQ board from Ninscope were utilized directly in this study. To alternately acquire the uniformly illuminated and structured-illumination images, the two LEDs were controlled by two current drivers modulated with gated PWM pulses to adjust their light power and synchronize with the exposure of the CMOS camera. The entire system was governed by a custom script developed in LabVIEW (National Instruments) via a DAQ card (USB6001, National Instruments). For imaging in freely moving mice, an open-source motorized commutator (***Kapanaiah and Kätzel, 2023***) was placed between the miniscope and the DAQ system.

### Data simulation

Simulation data (uniformly illuminated widefield images and structured illumination images) for the HiLo miniscope were generated using a method analogous to that employed in a prior study (***Zhou et al., 2018***). Both uniformly illuminated and structured illumination images comprise three components: in-focus neural activity (true signal), blurred out-of-focus neural activity (local background), and global background signal. In our simulations, in-focus and out-of-focus neurons were simplified as normalized Gaussian distributions with distinct dimensions and random spatial positions. The temporal dynamics of each neuron were introduced by multiplying the pixel values within the corresponding neuronal region by a real hippocampal neural activity dataset acquired via a widefield miniscope. A single frame from the identical dataset was directly adopted as the global background. The final widefield images were synthesized as a weighted sum of these three components, whereas the in-focus neurons for structured illumination images were multiplied by a one-dimensional periodic grid signal prior to summation with the other two components. Subsequent to this, the final HiLo images were reconstructed from the simulated widefield and structured illumination images using the identical algorithm implemented in the HiLo miniscope.

### Neural activity extraction

Widefield and HiLo images were downsampled from 480 × 480 to 240 × 240 pixels and motion-corrected using ImageJ (NIH, Bethesda, MD). For comparisons between widefield and HiLo imaging, neuronal signals were extracted using the CNMF-E (constrained non-negative matrix factorization for microendoscopic data) algorithm (***Zhou et al., 2018***). Because widefield and HiLo imaging differ in signal intensity and signal-to-noise characteristics, separate CNMF-E parameter sets were used (widefield: min_corr = 0.90, min_pnr = 8; HiLo: min_corr = 0.85, min_pnr = 3). For multi-plane imaging experiments, neuronal signals were extracted independently in each imaging plane using Suite2p (***Pachitariu et al., 2017***). Putative neurons were visually inspected based on the quality of their spatial footprints and calcium dynamics, and cells exhibiting unusually long decay kinetics or minimal activity were excluded from further analysis.

Spatial footprints identified by CNMF-E were treated as ROIs and applied to the corresponding widefield and HiLo images to extract fluorescence traces by averaging pixel intensities within each ROI. Baseline fluorescence (*F*_0_) was estimated dynamically using a sliding-window percentile method: for each time point, *F*_0_ was defined as the 10th percentile of fluorescence values within a ±15 s window centered on that time point. For time points near the beginning or end of a recording, the window was truncated to include only available data. Normalized fluorescence changes (Δ*F* ⁄*F*) were then computed as (*F* − *F*_0_)⁄*F*_0_. For matched-ROI comparisons, spatial footprints extracted from widefield data were applied identically to both widefield and HiLo recordings to ensure fair comparisons.

Three representations of neural activity were used for spatial information analyses: (1) ROI Δ*F* ⁄*F* traces; (2) raw fluorescence traces extracted by CNMF-E (*C*_raw_), with negative values set to zero; and (3) deconvolved calcium event traces estimated by CNMF-E (*S*), in which only events exceeding 3 standard deviations were retained.

To identify corresponding cells across imaging conditions (WF vs. HiLo or across focal planes), ROIs were matched using combined spatial and functional criteria. For each condition, ROIs were represented as binary spatial maps from which cell outlines were extracted, and the center of mass (COM) of each ROI was computed from its boundary coordinates. Spatial proximity was quantified as the Euclidean distance between COMs, and ROI pairs with center-to-center distances smaller than 15 *µ*m were considered candidate matches. Functional similarity was assessed by computing the Pearson correlation coefficient between calcium activity traces, with correlations greater than 0.5 indicating functional correspondence. ROIs were classified as representing the same cell only if they satisfied both spatial and functional criteria.

For ROI-based signal analyses, we used CNMF-E exclusively for ROI identification to ensure a standardized comparison across imaging conditions. However, because HiLo substantially improves image contrast (Figure 2G), ROI identification can potentially be achieved using simpler segmentation approaches, including Suite2p, Cellpose, or other image-based methods.

### SNR of calcium signals

To quantify the quality of extracted calcium signals, SNR was computed from the raw fluorescence traces. For CNMF-E-extracted signals, the raw temporal component (*C*_raw_) was used rather than the denoised trace (*C*). This choice was made because denoising procedures explicitly suppress noise fluctuations and alter the statistical properties of the signal, which can artificially inflate SNR estimates. Using *C*_raw_ preserves the residual measurement noise present after signal extraction and therefore provides a more direct assessment of signal quality. The SNR of calcium activity traces was computed using a peak-based approach designed to separate transient calcium events from baseline noise. For each fluorescence signal, candidate calcium transients were first detected using a peak-finding algorithm (findpeaks, MATLAB) with a minimum peak prominence set to three times the standard deviation of the entire signal. Detected peaks with durations shorter than 0.2 s were excluded to remove brief fluctuations unlikely to correspond to calcium events.

To estimate baseline noise, time windows surrounding each detected peak were removed from the signal. Specifically, for each peak, a window extending from 2 s before to 5 s after the peak time was excluded. For peaks occurring near the beginning or end of the recording, the exclusion window was truncated to remain within signal bounds. All excluded segments were removed, yielding a baseline signal composed of non-event activity. To further reduce contamination from residual calcium activity, baseline samples above the 80th percentile of the remaining signal distribution were discarded. The remaining values were treated as noise, and their standard deviation was used as the noise estimate.

The signal amplitude was defined as the mean of peak values exceeding the 90^th^ percentile of the detected peak distribution. The SNR was computed as the ratio between this mean peak amplitude and the standard deviation of the baseline noise. If no valid peaks were detected in a trace, the SNR was defined as zero.

### Multi-plane imaging

The electrowetting lens, or varioptic lens, is a flexible focusing device that modulates the shape of its liquid interface via voltage adjustment. Owing to the optical sectioning capability of the HiLo method, this lens is employed not only for achieving a consistent field of view (FOV) across multiple experiments but also for multi-plane imaging in our system. The relationship curve between the axial focal position and the applied voltage (***Figure 3—figure Supplement 1***A) was calibrated and subjected to linear fitting to guide subsequent multilayer optical sectioning imaging. In a 200 *µ*m-thick Thy1-GFP mouse brain slice, a high-contrast image stack with a maximum depth of 132 *µ*m (***Figure 3—figure Supplement 1***B) was acquired using the HiLo miniscope with a 2 V voltage step. For two-plane imaging (***Figure 3***B), the voltage of the electrowetting lens (EWL) was switched between two fixed values in synchronization with the acquisition of HiLo images. Given that the response time of the EWL exceeds the single-frame acquisition time of the HiLo miniscope, two frames acquired immediately after each voltage adjustment were discarded to avoid signal crosstalk between different imaging planes. The final multi-plane imaging acquisition rate achieved for two-plane *in vivo* imaging was 2.5 Hz.

### Open-field task

Recordings were conducted after animals had fully recovered from surgery. Prior to data collection, mice were habituated to the open-field arena for 2–3 days, during which they freely explored the arena under identical experimental conditions. Each session consisted of a single 30–40 min recording. Neural activity was recorded while mice freely explored a rectangular open-field arena (40 × 57 cm). Laminated A4 sheets displaying distinct visual patterns were affixed to each wall of the arena and served as distal spatial landmarks. To promote continuous and uniform exploration, small crumbs of sunflower seeds were periodically and randomly scattered throughout the arena. Animal position was tracked using an overhead near-infrared (NIR) camera. A NIR light source (940– 945 nm) mounted on the arena wall and within the camera’s field of view was used to synchronize behavioral tracking with calcium imaging.

### Animal position tracking

Animal position was extracted from video recordings of open-field behavior using a deep learning– based pose estimation pipeline. A total of 300 video frames (resolution: 1920 × 1080 pixels) were randomly sampled from recorded sessions and manually annotated using LabelMe (https://github.com/wkentaro/labelme). Three anatomical landmarks were labeled in each frame: the left ear, right ear, and body center. Annotated images were used to train an Ultralytics YOLO pose estimation model (https://github.com/ultralytics/ultralytics) using 10-fold cross-validation. Model training was performed for 200 epochs with a batch size of 16. Video reading and writing were accelerated using FFMPEGCV (https://github.com/chenxinfeng4/ffmpegcv), which provides efficient FFmpeg-based video processing optimized for deep-learning workflows. To further improve tracking robustness, frames that were difficult to predict—such as those containing curled body postures or partial occlusions (e.g., cables crossing the animal)—were manually selected and added to the training set for model fine-tuning. The model was subsequently fine-tuned for an additional 50 epochs with a batch size of 16. Predicted keypoint trajectories were temporally smoothed using a one-dimensional moving-average filter with a kernel size of 3.

### Place cell analysis

ROI Δ*F* ⁄*F* traces or CNMF-E–extracted signals were first aligned to the animal’s position using synchronization pulses recorded from the behavioral and imaging systems. Position samples corresponding to periods of immobility (speed ≤ 2.5 cm s^−1^) were excluded from subsequent analyses. The behavioral arena was divided into spatial bins (2.5 × 2.5 cm), and occupancy maps were calculated from the animal’s trajectory. Spatial bins visited for less than 0.2 s were excluded. For visualization, spatial tuning maps were generated by computing the mean neural activity within each spatial bin and smoothing the resulting maps with a Gaussian kernel (σ = 3.5 cm).

Spatial information content was calculated for each neuron as previously described (***Skaggs et al., 1996; Gauthier and Tank, 2018***):

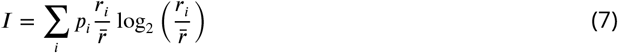

where *p*_i_ is the occupancy probability of bin _i_, *r*_i_ is the mean activity in bin _i_, and 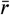 is the neuron’s mean activity across the entire session.

To assess statistical significance, spatial information values were compared to a shuffled distribution generated by circularly shifting each neuron’s activity trace relative to the behavioral trajectory (500 shuffles). Neurons were classified as place cells if their observed spatial information exceeded the 95^th^ percentile of the shuffled distribution.

### Bayesian decoding analysis

Animal position was decoded from population calcium activity using a Bayesian framework adapted from established spike-based decoders (***Zhang et al., 1998; Davidson et al., 2009; Lai et al., 2023; Ziv et al., 2013***). Neural activity was temporally aligned to behavioral tracking and binned at 333 ms (corresponding to five imaging frames). The posterior probability of the animal being at position *x* given population activity **r**(*t*) at time *t* was computed as

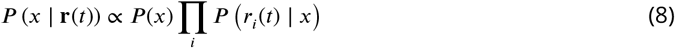

where *P* (*x*) is the prior probability of occupying bin *x*, and *P* (*r*_i_(*t*) ∣ *x*) is the likelihood of observing activity *r*_i_(*t*) from neuron _i_ at position *x*. Likelihood functions were modeled as Gaussian distributions, with means given by the spatial tuning curves *f*_i_(*x*) and variances estimated from the residual activity across time. A uniform prior over space was used to avoid bias toward highly occupied locations. Posterior probabilities were normalized across all spatial bins at each time point, and the decoded position was defined as the maximum a posteriori estimate.

To prevent circularity, decoding performance was evaluated using five-fold cross-validation with data split temporally into training and test sets. Spatial tuning curves were computed exclusively from the training data and applied to decode positions in the held-out test data. Decoding accuracy was quantified as the Euclidean distance between decoded and true positions at each time point. Overall decoding performance was summarized using the median decoding error across time.

### Closed-loop brain–machine interface

Head-fixed mice were presented with odors delivered by a vacuum-controlled airflow system and received sucrose rewards through a stainless-steel spout. Licking behavior was monitored using a contact-based lick detector coupled to the spout. An Arduino Uno board controlled odor presentation and reward delivery, and behavioral and task-related signals were collected using a USB data acquisition card (NI-USB-6501, National Instruments).

Mice were initially presented with a 1-s olfactory cue (citral, 5% v/v in propylene glycol) repeated 10 times, with inter-trial intervals randomly drawn from a uniform distribution between 15 and 25 s. The spatial footprints of all imaged neurons were extracted using Suite2p and applied to HiLo images to obtain fluorescence activity traces for each cell. Suite2p was used to enable rapid ROI extraction, minimizing delays associated with computationally intensive offline processing and thereby reducing the impact of neural drift during template generation. Neuronal responses were quantified by averaging activity between 0.5 and 1.5 s following odor onset and across the 10 presentations, yielding a population response vector (N neurons × 1) that served as the template.

After uploading the template, real-time similarity between the ongoing population activity and the template was computed as the Pearson correlation coefficient (*r*). Population activity vectors were updated at 15 Hz using activity averaged over the preceding 1-s window, producing a continuous estimate of representational similarity. Because baseline correlation values varied across animals, the reward threshold was manually set for each mouse.

During the acquisition block, a sucrose reward (5% w/v, ~5 µL) was delivered immediately when the real-time correlation exceeded the predefined threshold. Each reward trigger initiated a 5-s refractory period during which additional threshold crossings did not result in reward delivery. During the extinction block, threshold crossings were detected, but no rewards were delivered. Block durations were randomized across mice (acquisition: 18.2 ± 4.4 min; extinction: 10.0 ± 1.8 min; reacquisition: 8.33 ± 1.9 min; mean ± SEM).

The frequency of threshold-crossing events was calculated as the number of crossings occurring outside refractory periods divided by block duration. For analyses of pre- and peri-crossing lick rates, events occurring within 10 s of a preceding crossing were excluded to minimize contamination from consummatory licking.

## Acknowledgments

This work was supported by the Brain Science and Brain-like Intelligence Technology—National Science and Technology Major Project (2022ZD0207500), National Natural Science Foundation of China (32471089, 324B2032), CAMS Innovation Fund for Medical Sciences (CIFMS, 2024-I2M-ZD-013), and Chinese Institute for Brain Research (CIBR), Beijing. The funders had no role in study design, data collection, and interpretation, or the decision to submit the work for publication. The authors thank the Laboratory Animal Resource Center and the Optical Imaging Facility at CIBR for their services, and Jiayue Li at CIBR for assistance with figure artwork.

## Additional information

### Author contributions

Huixin Lin, Investigation, Formal analysis, Visualization, Methodology, Writing – original draft, Writing – review and editing; Shu Wang, Methodology, Investigation, Writing – review and editing; Yuhang Zhu, Investigation, Writing – review and editing; Zhaoyang Yin, Investigation, Writing – review and editing; Qingchun Guo, Conceptualization, Methodology, Supervision, Funding acquisition, Writing – original draft, Writing – review and editing; Jingfeng Zhou, Conceptualization, Methodology, Supervision, Funding acquisition, Writing – original draft, Writing – review and editing.

### Data and code availability

Source data and MATLAB scripts used in this study are available at OSF: https://osf.io/4659v. The OSF project has been assigned the DOI https://doi.org/10.17605/OSF.IO/4659V.

---

**Figure 1—figure supplement 1.**
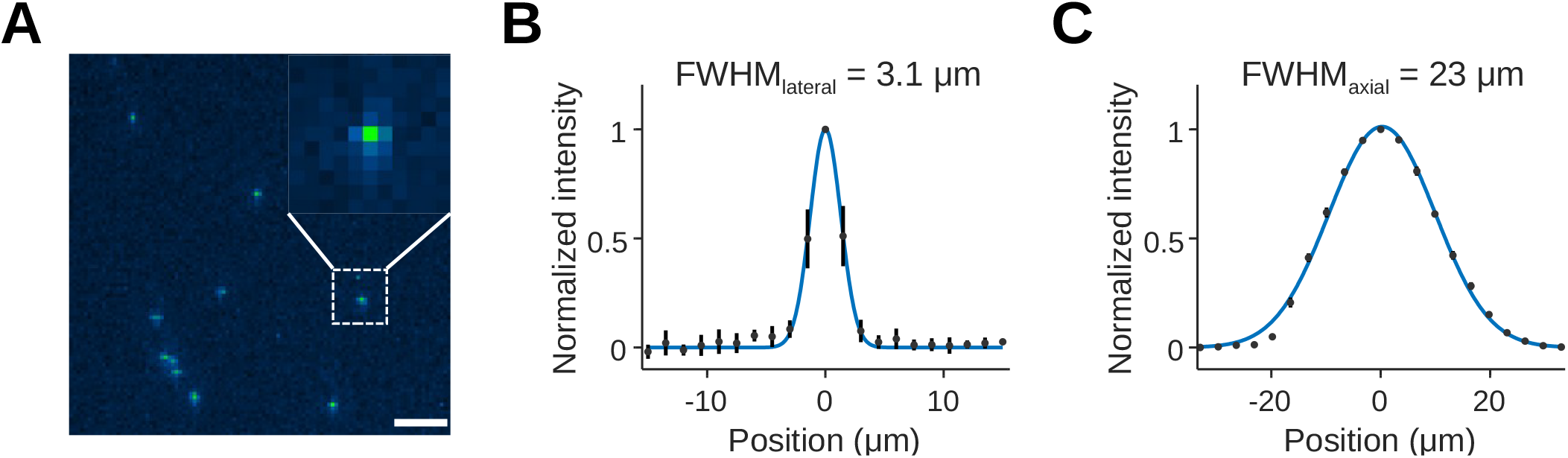
Lateral and axial resolution of the HiLo miniscope. (A) Representative image of sub-resolution fluorescent beads, 1 *µ*m in diameter, used to characterize optical resolution. The inset shows a magnified view of a single bead. Scale bar, 100 *µ*m. (B) Normalized lateral intensity profile along the x-axis, centered on the peak intensity of sub-resolution fluorescent beads. The Gaussian fit (blue line) yielded a lateral full width at half maximum (FWHM) of 3.1 *µ*m. Data are presented as mean ± s.d.; n = 8 beads. (C) Axial sectioning curve measured from a thin, uniform fluorescent plane as a function of defocus. The normalized axial intensity profile (black dots) and Gaussian fit (blue line) yielded an axial FWHM of 23 *µ*m.

**Figure 3—figure supplement 1.**
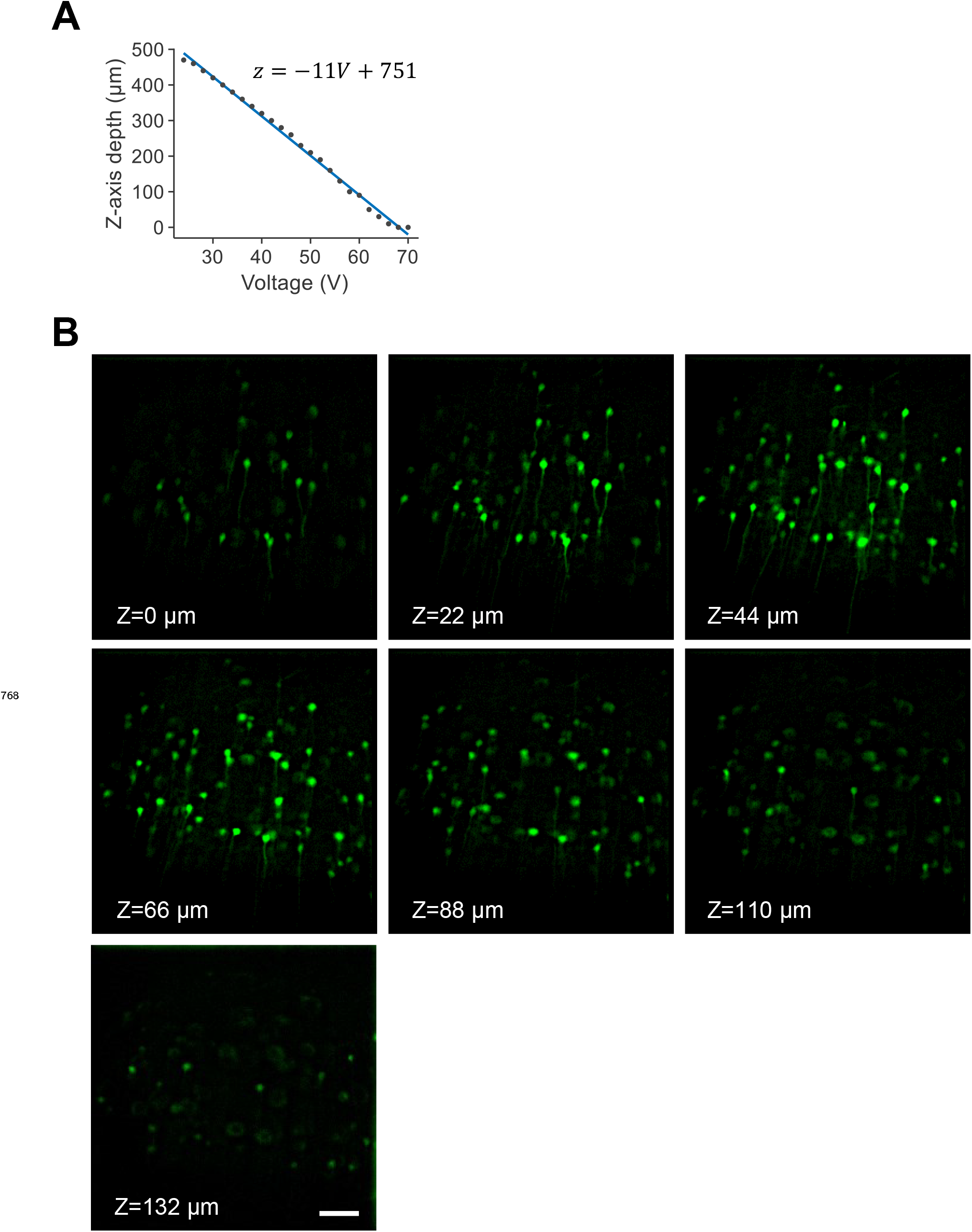
Multi-plane imaging with a liquid-lens–based HiLo miniscope. (A) Calibration of focal depth as a function of applied voltage for the electrowetting liquid lens. Linear fitting indicates reliable and repeatable axial displacement across the imaging range. (B) Representative HiLo images acquired at multiple focal depths, z = 0, 22, 44, 66, 88, 110, and 132 *µ*m, from a Thy1-GFP mouse brain slice. Scale bar, 100 *µ*m.

**Figure 4—figure supplement 1.**
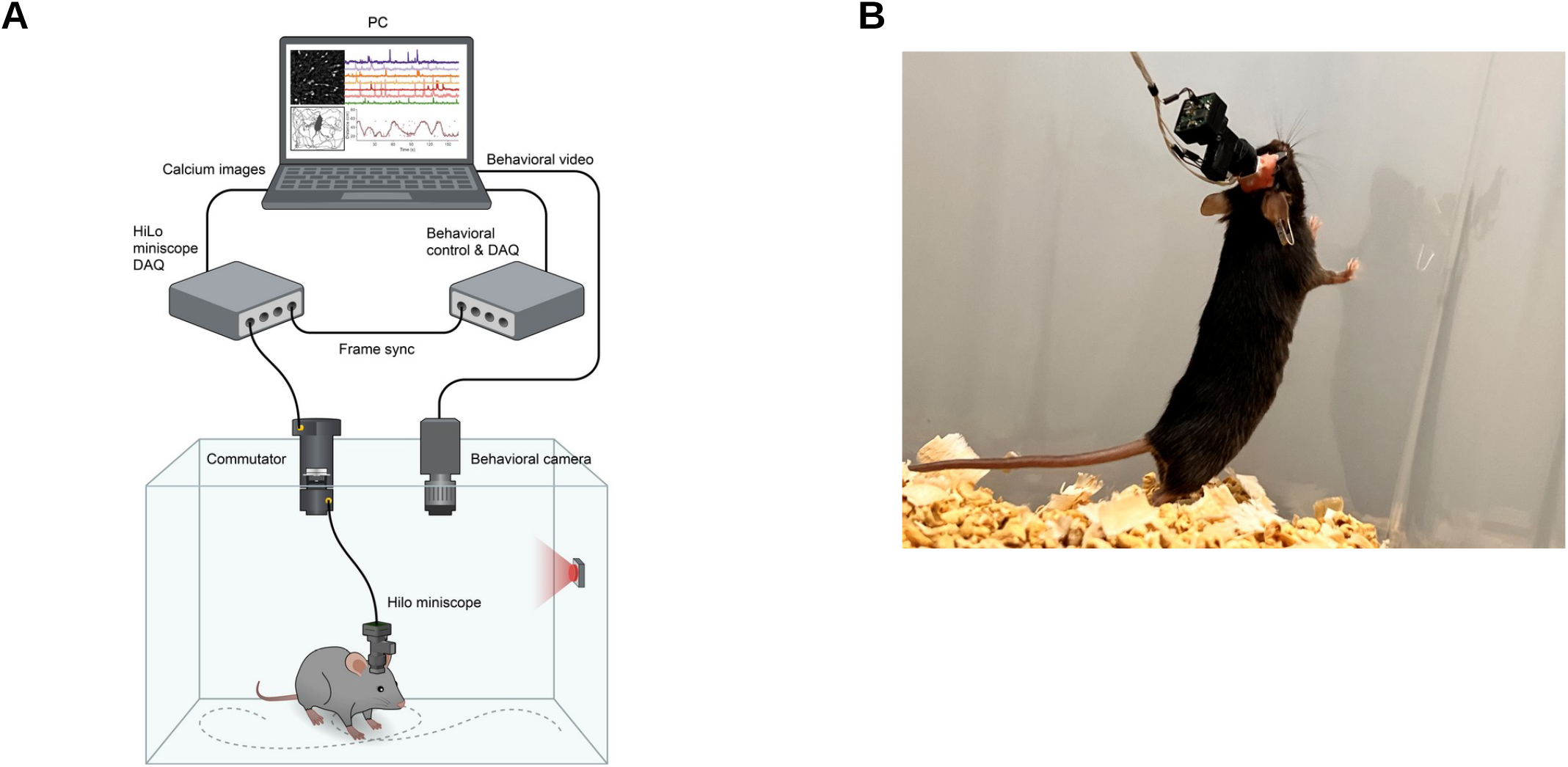
HiLo calcium imaging in an open-field arena. (A) Schematic of the freely moving imaging paradigm. Calcium imaging data were acquired using the HiLo miniscope while behavioral video was recorded simultaneously. A motorized commutator connected the miniscope to the DAQ device. Imaging and behavioral streams were synchronized and transmitted to a PC for online and offline analysis. (B) Photograph of a freely moving mouse implanted with a GRIN lens and wearing the HiLo miniscope during open-field exploration.

**Figure 4—figure supplement 2.**
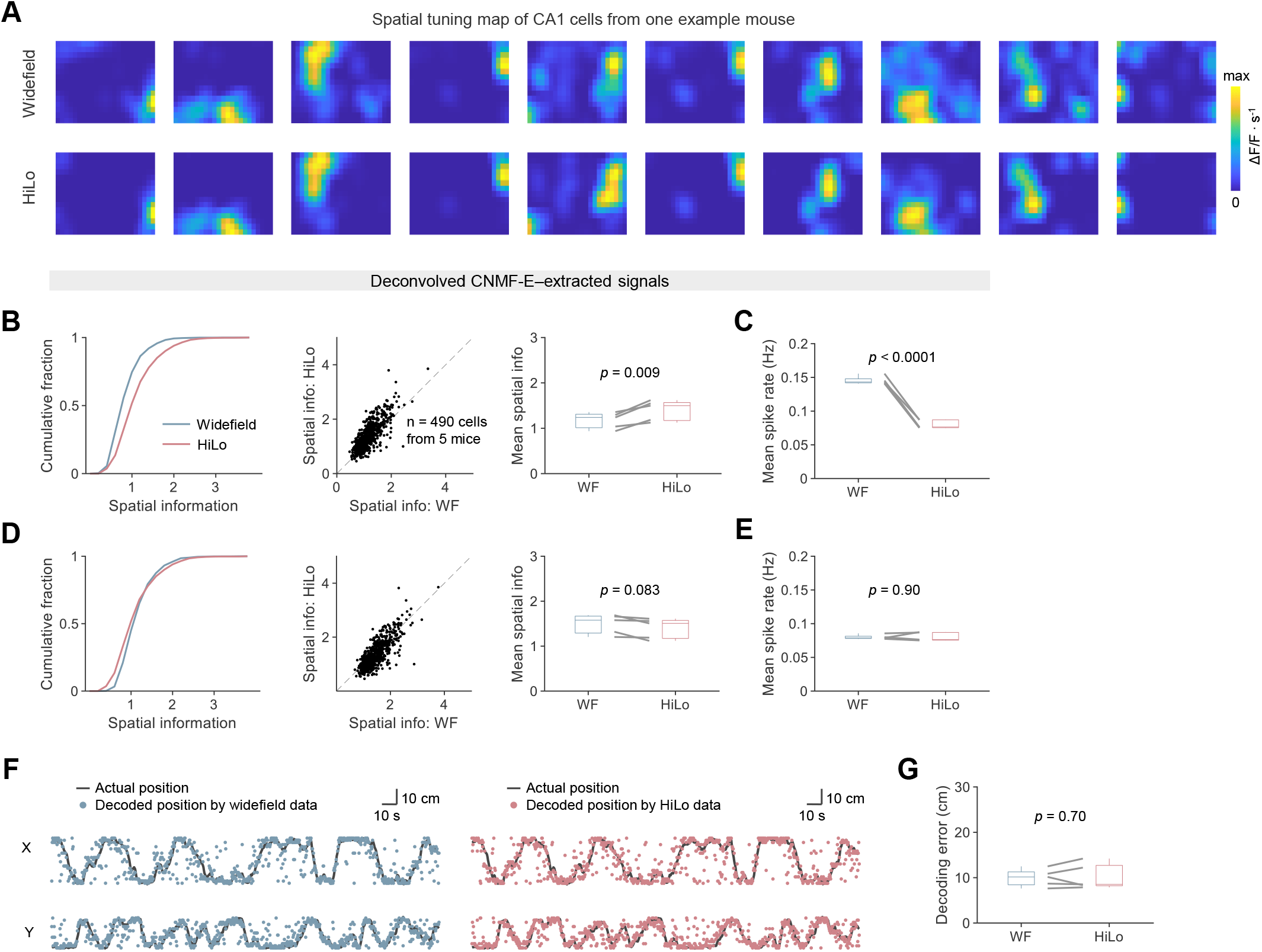
Spatial information readout from hippocampal CA1 during free exploration. (A) Representative spatial tuning maps of CA1 neurons from one example mouse, computed from widefield, top, and HiLo, bottom, imaging data. Maps show reduced background activity and sharper place fields in HiLo images. (B) Mean deconvolved spike rate. (C) Spatial information computed from deconvolved CNMF-E–extracted fluorescent signals. Left, cumulative distributions of spatial information for widefield, blue, and HiLo, red, data. Middle, scatter plot comparing spatial information values for matched neurons, n = 490 cells. Right, summary of mean spatial information for matched cells across imaging modalities. (D) Mean deconvolved spike rate after matching event rates by randomly removing a subset of widefield-estimated spike events. (E) Spatial information calculated from spike-rate-matched CNMF-E–extracted deconvolved activity, displayed in the same format as in (C). (F) Bayesian decoding of animal position using deconvolved CNMF-E–extracted signals. Actual trajectories are shown in black, with decoded positions overlaid as colored dots for widefield, left, and HiLo, right, data. (G) Decoding performance quantified as median decoding error for widefield and HiLo data. In (B)–(E) and (G), paired lines indicate within-session comparisons; boxes show median and interquartile range; whiskers indicate extreme values; statistical significance was assessed using paired *t*-tests.

**Figure 5—figure supplement 1.**
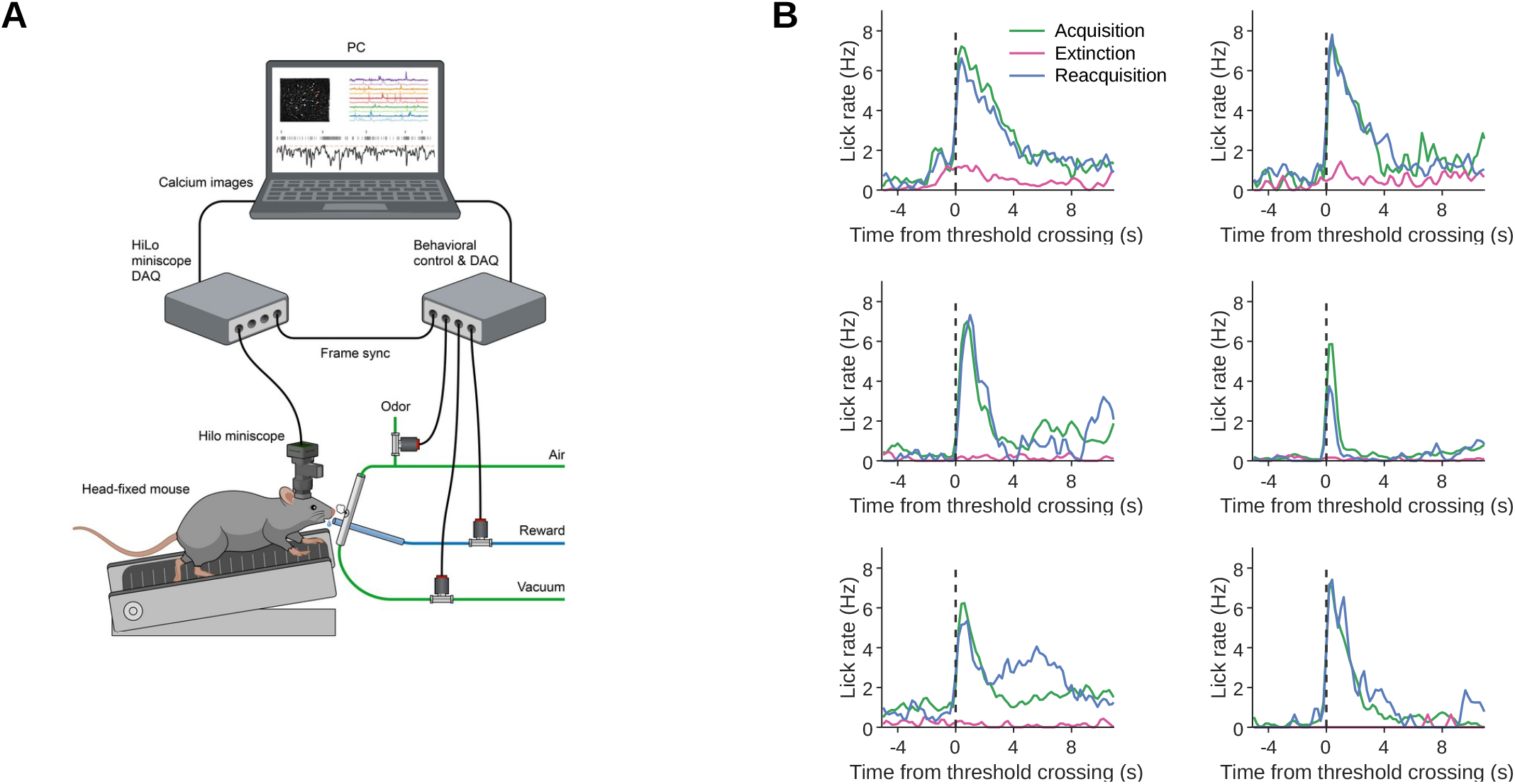
Closed-loop calcium imaging and lick behavior during BMI training. (A) Schematic of the experimental setup for head-fixed mice performing a closed-loop brain– machine interface task. The HiLo miniscope records calcium signals from neurons. Behavioral events, including odor delivery, reward delivery, and vacuum, are controlled by a separate DAQ system and synchronized with imaging frames. The system allows real-time monitoring and integration of neural activity and behavioral outputs. (B) Lick rate as a function of time relative to threshold-crossing events, dashed line at 0 s, for all six mice.

